# Connectivity Patterns in Cognitive Control Networks Predict Naturalistic Multitasking Ability

**DOI:** 10.1101/296475

**Authors:** Tanya Wen, De-Cyuan Liu, Shulan Hsieh

**Affiliations:** MRC Cognition and Brain Sciences Unit, University of Cambridge, United Kingdom; Department of Psychology, Asia University, Taichung, Taiwan; Department of Psychology; Institute of Allied Health Sciences; Department of Public Health, National Cheng Kung University, Tainan, Taiwan

**Keywords:** multitasking, functional connectivity, prefrontal networks, hierarchical organization, multivariate pattern analysis

## Abstract

Multitasking is a fundamental aspect of everyday life activities. To achieve a complex, multi-component goal, the tasks must be subdivided into sub-tasks and component steps, a critical function of prefrontal networks. The prefrontal cortex is considered to be organized in a cascade of executive processes from the sensorimotor to anterior prefrontal cortex, which includes execution of specific goal-directed action, to encoding and maintaining task rules, and finally monitoring distal goals. In the current study, we used a virtual multitasking paradigm to tap into real-world performance and relate it to each individual’s resting-state functional connectivity in fMRI. While did not find any correlation between global connectivity of any of the major networks with multitasking ability, global connectivity of the lateral prefrontal cortex (LPFC) was predictive of EVET score. Further analysis showed that multivariate connectivity patterns within the sensorimotor network (SMN), and between-network connectivity of the frontopartietal network (FPN) and dorsal attention network (DAN), predicted individual multitasking ability and could be generalized to novel individuals. Together, these results support previous research that prefrontal networks underlie multitasking abilities and show that connectivity patterns in the cascade of prefrontal networks may explain individual differences in performance.

## 1. Introduction

Multitasking is a fundamental aspect of everyday activities. The ability to multitask closely relates to flexible cognitive control (Rothbart & Posner, 2015), in particular, the selection and monitoring of higher order internal goals while other sub-goals are being performed. This is subserved by a set of frontal and parietal regions, which together assemble the required cognitive operations for task-relevant behavior (Norman & Shallice, 1986; Duncan & Owen, 2000; Miller & Cohen, 2001; Duncan, 2013; Cole & Schneider, 2007; Fedorenko et al., 2013).

This set of fronto-parietal regions, collectively known as the cognitive control network, shows especially high global connectivity, allowing information to be coordinated throughout the brain (Cole et al., 2010). Resting-state global connectivity in regions of this network, especially the lateral prefrontal cortex (LPFC), has been found to be correlated with attention (Rosenberg et al., 2015), cognitive control capacity (Cole et al., 2012), and general fluid intelligence (Song et al., 2008; Cole et al., 2012; Hearne et al., 2016). Together, these evidence suggest that functional connectivity in networks related to cognitive control may be the key underlying individual differences in carrying out complex task demands.

Previous studies linking connectivity to cognitive ability have employed neuropsychological assessments (Cole et al., 2012) such as the Raven Advanced Progressive Matrices (Raven et al.,1998) and the Cattell Culture Fair Test (Cattell, 1967), or conventional laboratory tasks with a limited number of experimentally controlled manipulations (Chen et al., 2016; Cole et al., 2013; Dux et al., 2009; Garner & Dux, 2015). These tasks often involve responses to shapes or numbers on the screen according to a set of task rules. While they provide useful insight into the neural underpinnings of general executive function, they measure separable cognitive abilities compared to multitasking (Kievit et al., 2014). In fact, several keystone studies have found that patients with frontal lobe damage show deficits in everyday multitasking (such as planning a dinner or doing grocery shopping), even though some had superior IQ and intact perfomances on neuropsychological tests of attention, memory, and executive functions (Shallice & Burgess, 1991; Burgess et al., 2000; Roca et al., 2011). This suggests multitasking requires an aspect of executive function that is not captured by standardized tests (Manly et al., 2002; Roca et al., 2009, 2011, 2012).

The current study examined the pattern of several well-defined functional networks (Power et al., 2011) during rest and performance on a simulated virtual real-life task, the Edinburgh Virtual Errands Test (EVET, Logie et al., 2011; Trawley et al., 2011, 2013). This test builds upon the Multiple Errends Test (Shallice & Burgess, 1991) to assess multitasking ability in the healthy population. We related each individual’s score of multitasking performance to resting-state functional connectivity. We focused on the networks involving the frontal, parietal, and sensorimotor regions, as these regions are thought to be involved in the neural architecture supporting cognitive control and goal-directed behavior in external tasks (Koechlin et al., 2003; Dixon et al., 2017; Miller & Cohen, 2001). These networks included the frontoparietal network (FPN), dorsal attention network (DAN), and sensorimotor network (SMN). However, in order to not overlook any other potential networks that may support multitasking, we included in our analysis other major functional networks, together covering most of the brain, for completeness (Power et al., 2011; Hearne et al., 2016). In the first analysis, we applied a linear regression analysis to examine whether global connectivity in cognitive control networks predicts multitask performance indexed by EVET score. Additionally, as the LPFC has been implicated as a critical region in cognitive function in previous studies (Koechlin et al., 1999; Song et al., 2008; Cole et al., 2012; 2015), we additionally analyzed global connectivity of the LPFC region and related it to EVET performance. However, global connectivity is a relatively coarse index as it sums all the functional connections from a network, creating an average summary index. Multivariate analysis takes account the ‘pattern’ of functional connections rather than the overall ‘strength’ (Kriegeskorte & Bandettini, 2007; Mur et al., 2009). Therefore, in our second analysis, we applied a support vector machine regression (SVR) model and leave-one-subject-out cross-validation to predict EVET scores of individual subjects (Dosenbach et al., 2010). We hypothesize that intrinsic functional connectivity within and between networks associated with cognitive control should be predictive of individual multitasking performance.

## 2. Methods

### 2.1 Participants

106 healthy volunteers (57 females, mean age 22.85, SD = 2.04) were included in the final analyses of the study. Participants were recruited via online advertisements. An additional 11 participants were excluded (five had excessive motion with mean frame displacement more than two standard deviations away from the group mean, one with abnormal cerebellar structure, and five who did not complete the full experiment). Inclusion criteria were right-handedness (indexed by the Edinburgh Handedness Inventory); no depression or anxiety (Beck Depression Inventory score < 14, Beck Anxiety Inventory score < 7); and no sleep disorders, history of drug use, chronic disease, mental health or neurological issues. Furthermore, none of the included participant’s head motion exceeded 2 mm / 2 degrees across the whole scanning run. Procedures were carried out in accordance with ethical approval obtained from the National Cheng Kung University Research Ethics Committee, and participants provided written, informed consent before the start of the experiment.

### 2.2 Experimental procedures

Inside the MRI scanner, participants were asked to relax with their eyes closed. Participants were instructed to keep their head still, remain awake, and not to think of any one persistent thought throughout the scan. First, a structural image was acquired (~3.6 minutes), followed by resting-state functional EPI images (~ 8 minutes).

For behavioral measures, we used the EVET (Logie et al., 2011; Trawley et al., 2011; Trawley et al., 2013; http://www.psy.ed.ac.uk/resgroup/MT/index4.html) to evaluate their multitasking ability. This was usually performed within a weeks’ time frame relative to the scan. EVET requires participants to complete eight errand tasks efficiently within 8 minutes while navigating through a simulated environment on a computer. The environment consisted of a four-story building with a set of stairs and five rooms along the left and right ends of each floor surrounding a central elevator (see Figure 1 for sample screen shots). Example errands include “pick up brown package in (room) T4 and take to (room) G6”, “meet person S10 before 3:00 min”, “sort as many red and blue binders as you can in room S2”. The set of diverse subtasks taps into multiple cognitive functions including planning, retrospective and prospective memory, visuospatial and verbal working memory, sustained attention, and task switching. The test was designed to evaluate participants’ ability to complete a complex task with higher ecological validity than conventional experimental manipulations.

**Figure 1.**
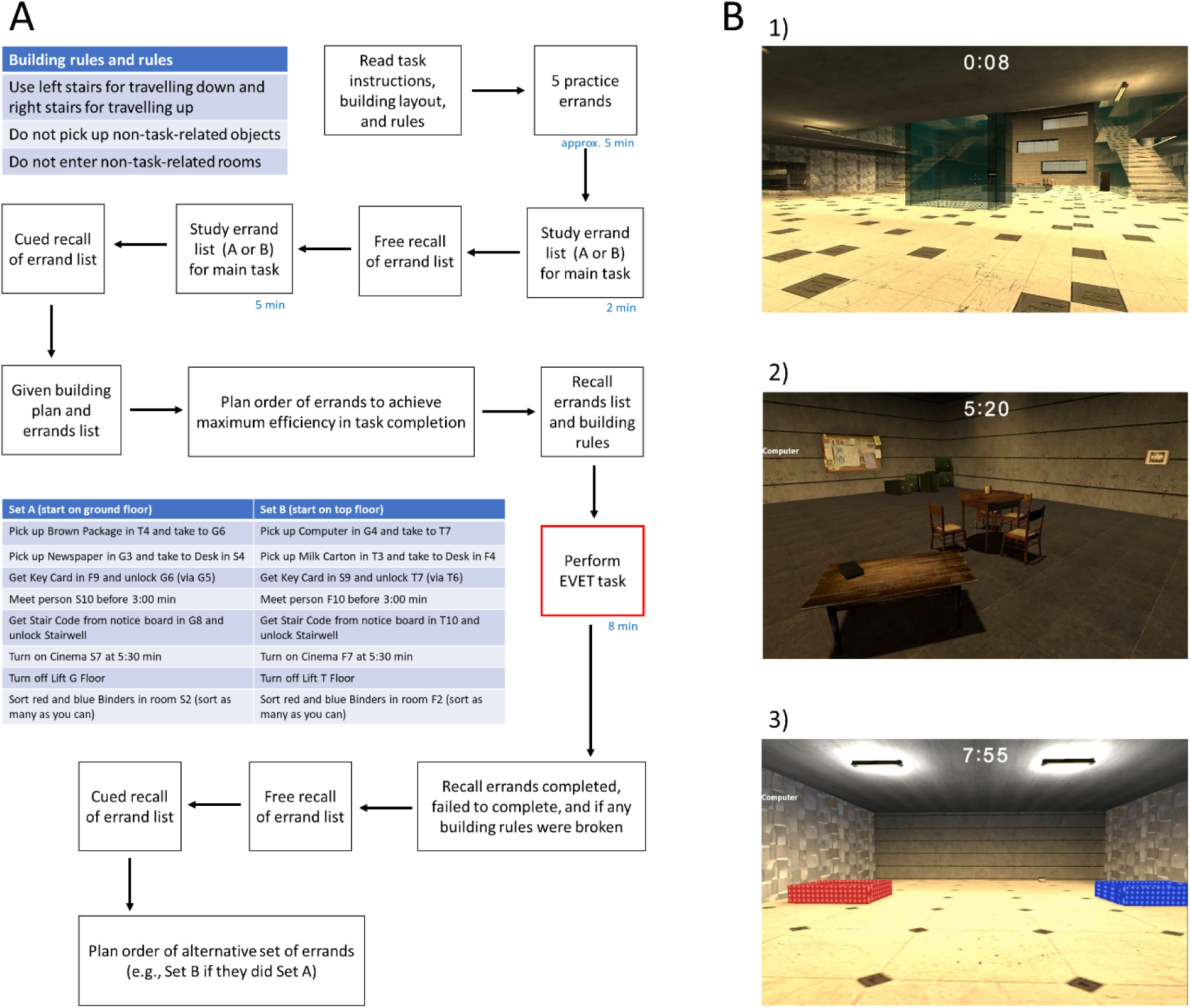
A. Pipeline of Edinburgh Virtual Errands Test (EVET) experiment. Participants were well familiarized with the building plan and task errands and performed the EVET for 8 minutes. B. Screen shots of the EVET environment. 1). Lobby with the elevator in the middle and staircases on the sides. 2). Room F9 where participants need to pick up a key card to unlock room G6. 3). Room F2 where participants need to sort as many red and blue binders as possible.

The execution of the EVET followed Logie et al. (2011). The EVET consists of two sets of tasks each with eight errands; 54 participants performed set A and 52 performed set B. First, participants were also informed of a building rule that requires to use the left stairs for travelling down and right stairs for travelling up; and the task rules of not entering irrelevant rooms or picking up non-task-related objects. Before the main experiment, participants practiced using the keyboard and mouse controls to move around the building and complete five practice errands. In the main EVET experiment, participants were given 2 minutes to study their errand list, followed by free recall, then another 5 minutes of further study and a test of cued recall. Then, participants were asked to plan the order in which they should perform each errand to achieve maximum efficiency in task completion. Next, they were asked to verbally recall the errand list and building rules until they could recall 100% of the list. Participants then performed the EVET for 8 minutes. Afterwards, they were asked to recall the errands they had attempted or failed to complete. Participants were cued about any errands they had omitted in the posttest recall. Finally, participants were given the alternative set of errands and were asked to plan the order of errands (e.g., participants who performed set A were asked to plan set B), which provided another measure of planning, but without performing the EVET a second time. Finally, a general “EVET score” was calculated based on participants’ overall performance (accounting for completed errands and incorrect actions). Points were added for each errand completion, bonus points were awarded based on number of folders sorted, time discrepancy for timed errands; and points were deducted for picking up incorrect objects, entering rooms not on the errand list, and breaking the building rules. Bonus and penalty points were given/deducted on a five-point-scale (0-4) based on a cutoff score that was calculated based on the frequency distribution of raw scores among the participant sample in Logie et al. (2011). Therefore, the minimum possible score was −12 and the maximum 20.

### 2.3 Imaging parameters and data analysis

Imaging was performed using a GE MR750 3T scanner (GE Health care, Waukesha, WI, USA) at the National Cheng Kung University Mind Research and Imaging Center MRI center. High-resolution anatomical T1 images were acquired using fast-SPGR, consisting of 166 axial slices (TR = 7.6 ms, TE = 3.3 ms, flip angle = 12◦, 224 × 224 matrices, slice thickness =1 mm), which lasted 218 s. Functional images were acquired with a gradient-echo echo-planar imaging (EPI) pulse sequence (TR = 2000 ms, TE = 30 ms, flip angle = 77◦, 64 × 64 matrices, FOV = 22 × 22 cm^2^, slice thickness = 4 mm, no gap, voxel size = 3.4375 mm × 3.4375 mm × 4 mm, 32 axial slices covering the entire brain). A total of 245 volumes were acquired; the first five served as dummy scans and were discarded to avoid T1 equilibrium effects.

Data were preprocessed using SPM 8 (http://www.fil.ion.ucl.ac.uk/spm) and the Data Processing & Analysis for Brain Imaging toolbox (DPABI, Yan et al., 2016) implemented in Matlab (The MathWorks, Inc., Natick, MA, USA). EPI images were slice-time corrected and realigned to correct for head motion using rigid-body transformation. The T1 image was coregistered to the mean EPI image and was normalized to the MNI template. The normalization parameters of the T1 image were applied to all functional volumes. Nuisance time series (motion parameters, ventricle and white matter signals) were regressed out. The functional data were spatially smoothed with a 6 mm Gaussian kernel. Finally, images were band-pass filtered at 0.01–0.08 Hz to remove scanner drift and high frequency noise (e.g., respiratory and cardiac activity). There was no correlation between EVET score and mean frame displacement (*r* = 0.19, *p* = 0.06). Furthermore, correcting for micro-head-motion (removing volumes with FD > 0.2 mm) did not alter the results of the study.

### 2.4 Functional connectivity analysis

The analysis pipeline is illustrated in Figure 2. Large-scale functional networks were identified based on an atlas that contained 264 nodes of interest (Power et al., 2011). The 10 major networks included the frontoparietal network (FPN), cingulo-opercular network (CON), default mode network (DMN), salience network (SN), dorsal attention network (DAN), ventral attention network (VAN), visual, auditory, sensorimotor networks (SMN), as well as subcortical regions. For each individual, mean activity was extracted from the 264 nodes of interest (10 mm sphere), and functional connectivity was estimated between each pair of nodes with Pearson’s correlation and were Fisher z-transformed. For univariate analysis, we calculated the average connectivity of each node with the rest of the nodes in the brain, which has been defined as global brain connectivity, or unthresholded weighted degree centrality (Cole, Anticevic, Repovs, & Barch, 2011; Cole et al., 2010; Cole et al., 2012; Rubinov & Sporns, 2011). Global brain connectivity was averaged within each functional network, and this served as a summary statistic of the network’s global connectivity. For multivariate analysis, the assemble of connections of each node in the network with all other nodes in the brain served as the feature vector for that network. All multiple comparisons from statistical tests were Sidak-corrected for multiple comparisons (10 networks and 55 possible within- and between-network connections).

**Figure 2.**
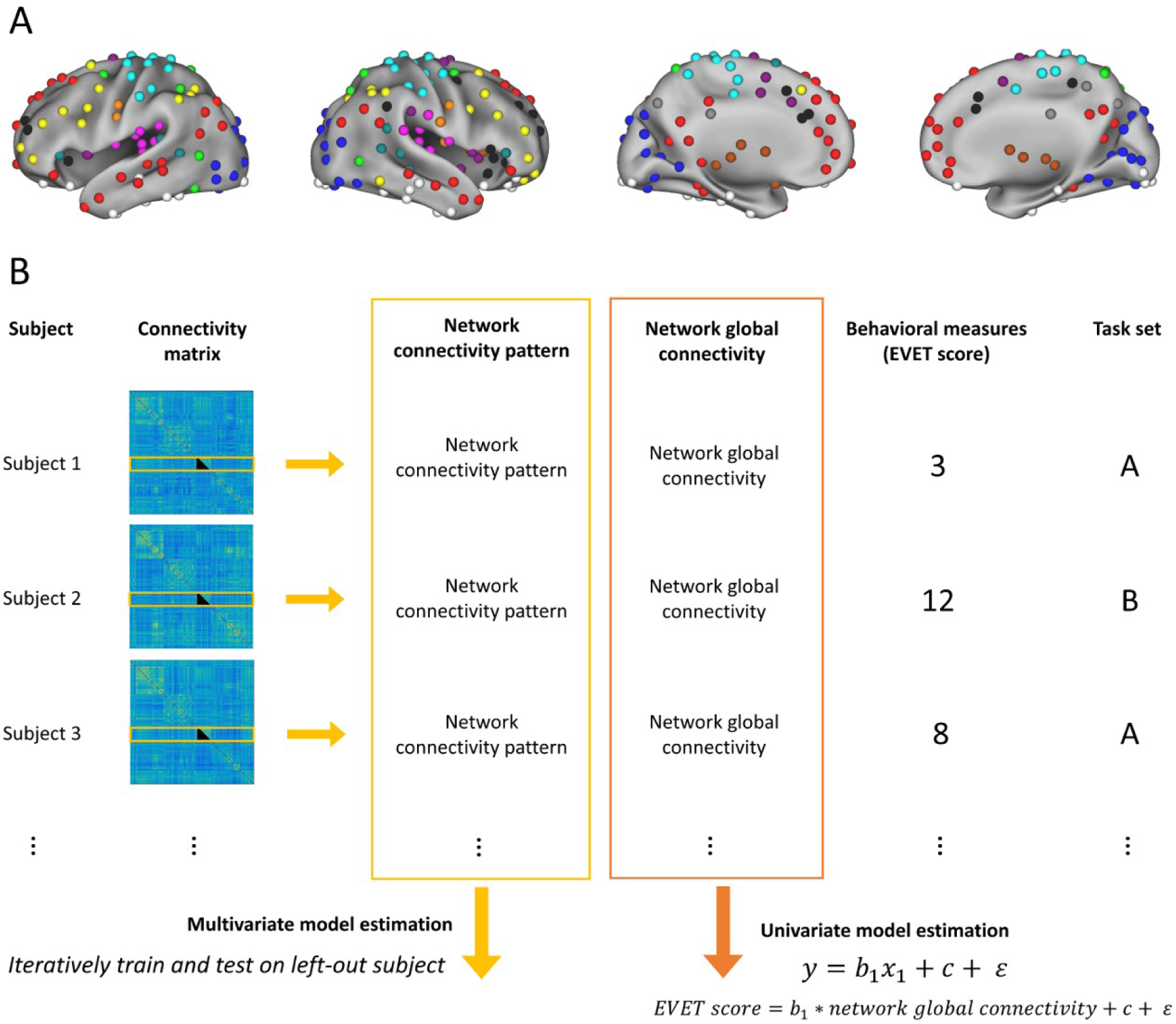
Analysis pipeline. A. The 264 nodes used in the experiment. Each color represents a distinct network. The figure was adapted from Power et al. (2011). B. A functional connectivity matrix was created for each participant. For each network, the connectivity of a node with the rest of the brain was extracted. A functional connectivity matrix was created for each participant. For each network, the connectivity of a node with the rest of the brain was extracted. In the univariate analysis, the average of these connections within a network was defined as the network global connectivity index, which was used in a linear regression with which task set the participant performed, to predict the participants’ EVET scores. In the multivariate analysis, within a network, all connections of each of the networks’ nodes with the remaining nodes in the brain served as a pattern vector that was included in a support vector machine regression (SVR) model. For each iteration, one participant was left out before training the SVR, and functional connectivity data from the left-out participant was used to predict their EVET score.

### 2.5 Univariate analysis

#### 2.5.1 Network global connectivity

For each network, a linear regression model was used to estimate participants’ EVET scores according to the network’s global connectivity. The linear model was as follows:

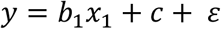

where y is the EVET score, x_1_ is network’s global connectivity measure, c is the intercept, *ε* is the error term, and b_1_ represents the main effect of the relationship between network global connectivity and EVET scores.

Furthermore, we subdivided each network’s global connectivity into mean within- and between-network connectivity to see whether relationships between specific networks are related to individuals’ multitasking performance. The same linear regression was performed on each network-network connectivity measure.

#### 2.5.2 ROI global connectivity

To compare with previous studies that found global connectivity in frontoparietal regions predictive of cognitive function (e.g., Song et al., 2008; Cole et al., 2012), we defined a LPFC node in the Power et al. (2012) template as the node closest to the LPFC ROI in Cole et al. (2012; MNI coordinates x = −44.2, y = 13.7, z = 29.8). The critical node had coordinates x = −47, y = 11, z = 23, and was found to be located in the FPN. Gobal connectivity of this LPFC node was defined as the mean functional connectivity of the LPFC with the other 263 nodes in the brain. We performed a linear regression as above, to estimate participants’ EVET scores from global connectivity in LPFC.

### 2.6 Multivariate analysis

In addition to average network connectivity, patterns of connectivity might be important in cognitive functioning. We calculated for each node the functional connectivity between that node and the remaining nodes in the brain. For each network, all functional connections of the nodes it consists of served as a pattern vector input of the model. To determine whether the pattern of network functional connectivity predicted multitasking ability in novel individuals, a leave-one-subject-out cross-validation was employed. In each set of n-1 individuals an SVR model was trained using each network’s connectivity patterns and the n-1 EVET scores, to predict the EVET score of the left out individual. Pearson’s correlation of predicted and actual EVET scores was used to assess the model’s predictive power.

Furthermore, to understand which functional connections are important contributing features in networks that showed above-chance decoding, functional connectivity patterns were further subdivided into within-network and between-network connectivity. For between-network connectivity, we separated each network’s functional connectivity with the 9 other networks in the brain. The same multivariate analysis was repeated with network–network connectivity patterns. Pearson’s correlation between participants’ predicted and actual EVET scores was used to evaluate each model’s predictive power.

In addition to leave-one-out cross-validation, we repeated our analyses using k-fold cross-validation, with k being 5 and 10. The results remained the same.

## 3 Results

### 3.1 Behavioral performance

The EVET scores ranged from −1 to 19 in our sample. Participants that did task set A (*mean* = 10.61, *SD* = 5.74) did not show any significant differences in performance from those that did set B (*mean* = 11.35, *SD* = 5.34; *t* = − 0.68, *p* = 0.50, two-tailed). There were significant correlations of the EVET score with initial recall of the task list (r = 0.41, p < 0.0001), pre-task planning (r = 0.38, p < 0.0001), following of their plan during the task (r = 0.68, p < 0.0001), recount of performance (r = 0.66, p < 0.0001), task list recall after EVET execution (r = 0.32, p < 0.001), and planning of the alternative task (r = 0.23, p = 0.02). There were no correlations between EVET performance and BAI (*r* = 0.01, *p* = 0.94) or BDI (*r* = 0.02, *p* = 0.84). We therefore only focused on the general EVET score in the fMRI analysis.

### 3.2 Relationship between network global connectivity and multitasking performance

#### 3.2.1 Network global connectivity

We examined the linear regression of EVET scores predicted by network global functional connectivity and task set. None of the 10 networks showed significant global connectivity and multitasking relationships after Sidak correction. Among the networks tested, the highest global connectivity coeffient was for the FPN (*b1* = 6.72, *SE* = 0.87, *p* = 0.09 (uncorrected), *p* = 0.59 (corrected), JZS Bayes Factor = 4.11).

Furthermore, we repeated the analysis subdividing into within- and between-newtork mean functional connections. However, none of the network-network connectivity measures showed any significant relationship between EVET scores after correcting for multiple comparisons. The highest regression coefficient was from the mean FPN-DAN connectivity [*b1* = 8.02, *SE* = 3.27; p = 0.02 (uncorrected), p = 0.58 (corrected), JZS Bayes Factor = 16.22].

#### 3.2.2 ROI global connectivity

Figure 3 illustrates the relationship between LPFC global connectivity and EVET scores. Linear regression of LPFC global connectivity was predictive of EVET scores (*b1* = 6.38, *SE* = 3.14, *p* = 0.04). This reconciles with previous studies showing that intrinsic connectivity of regions within the cognitive control network is related to cognitive function.

**Figure 3.**
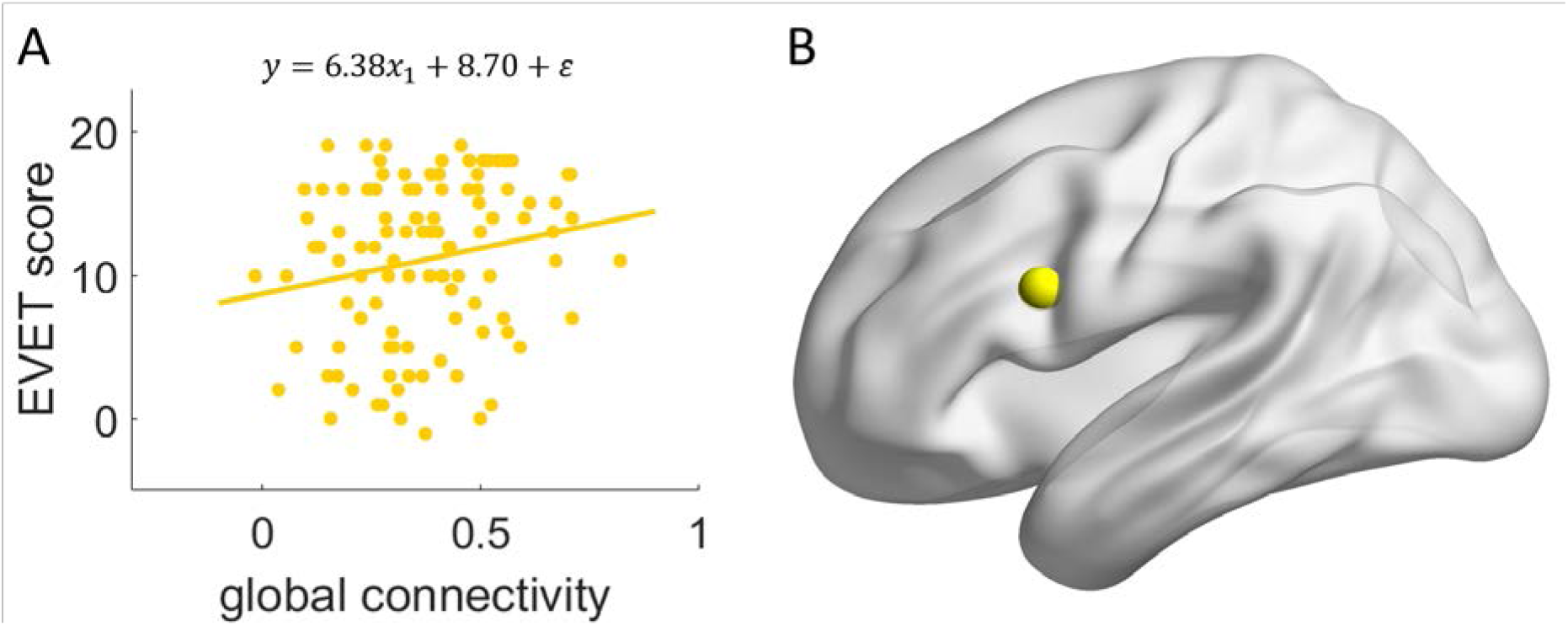
A is a scatter plot showing the relationship between global connecitvity of the LPFC with the remaining nodes in the brain is predictive of EVET performance. B illustrates the location of the LPFC node used in the analysis.

### 3.3 Predicting individual subject multitasking ability from network functional connectivity

Figure 4 illustrates each participant’s observed EVET score plotted against their predicted EVET score from the SVR. Functional connectivity patterns of the SMN, FPN, and DAN predicted multitasking performance in novel individuals, with the observed and predicted scores significantly correlated [SMN: *r* = 0.35, *p* < 0.001 (uncorrected), *p* < 0.01 (corrected); FPN: *r* = 0.30, *p* < 0.005 (uncorrected), *p* = 0.02 (corrected); DAN: *r* = 0.35, *p* < 0.001 (uncorrected), *p* < 0.01 (corrected)]. No other network showed significant cross-validation [all *rs* < 0.14, all *ps* > 0.14 (uncorrected)].

**Figure 4.**
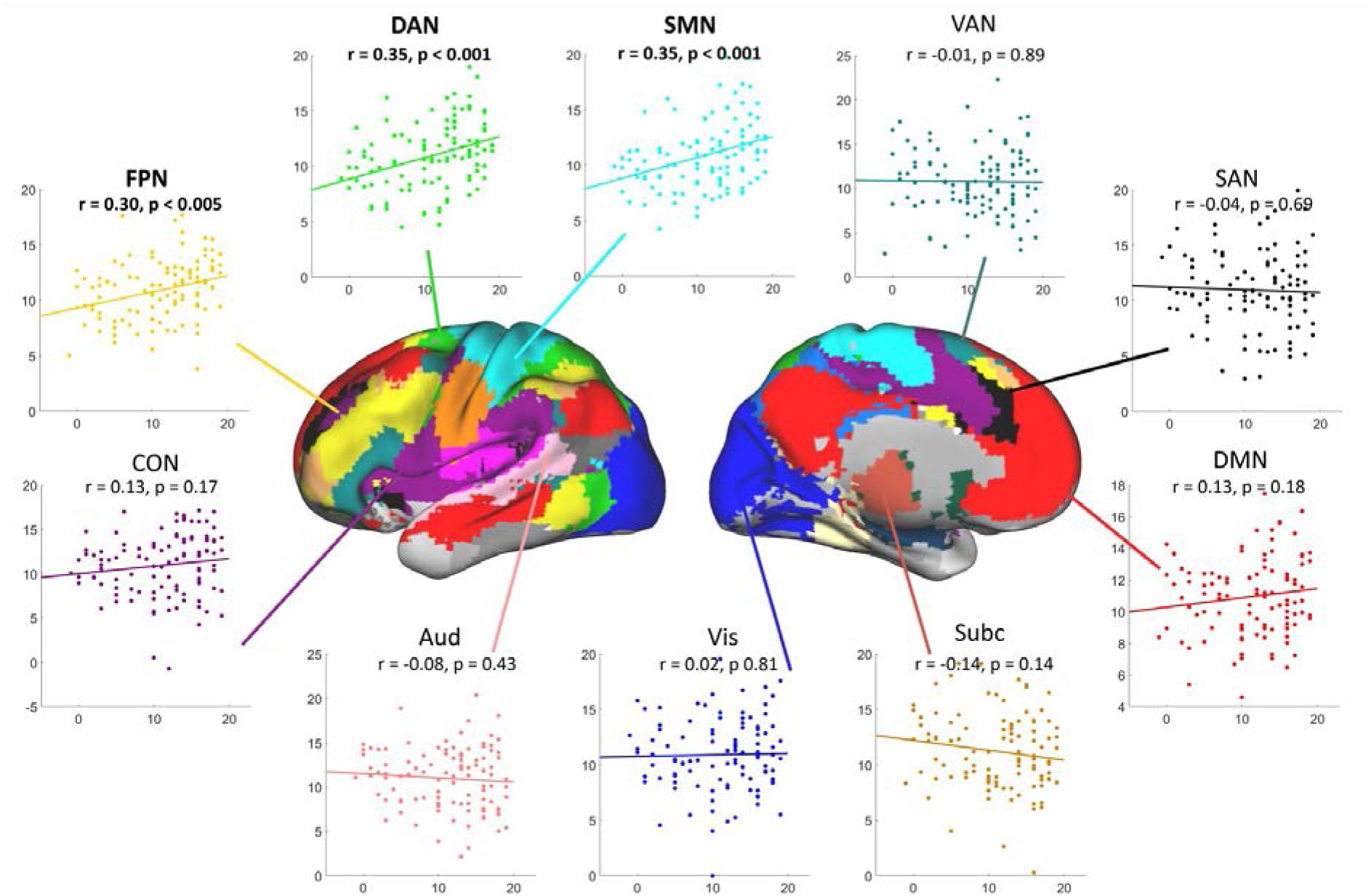
Multivariate pattern analysis results using network connectivity patterns. Scatter plot shows the correlation of observed EVET scores (x axis) with predicted EVET scores (y axis). Regions with connectivity patterns that showed significant decoding are labeled in bold and include the anterior-posterior gradient spanning the FPN, DAN, and SMN. The brain parcellation in the center was adapted from Power et al. (2011) and shows the color-coded functional networks.

Figure 5 illustrates the predictive power of each of the significant network–network functional connectivity patterns. Within-network connectivity of the SMN [*r* = 0.40, *p* < 0.001 (uncorrected), *p* = 0.001 (corrected)] and between-network FPN-DAN connectivity [*r* = 0.34, *p* < 0.001, *p* = 0.02 (corrected)] showed above chance prediction of novel individuals’ EVET score. No other connectivity patterns outside these three networks were predictive of individuals’ multitasking ability [all *r*s < 0.31, all *p*s > 0.001 (uncorrected), all *p*s > 0.06 (corrected)].

**Figure 5.**
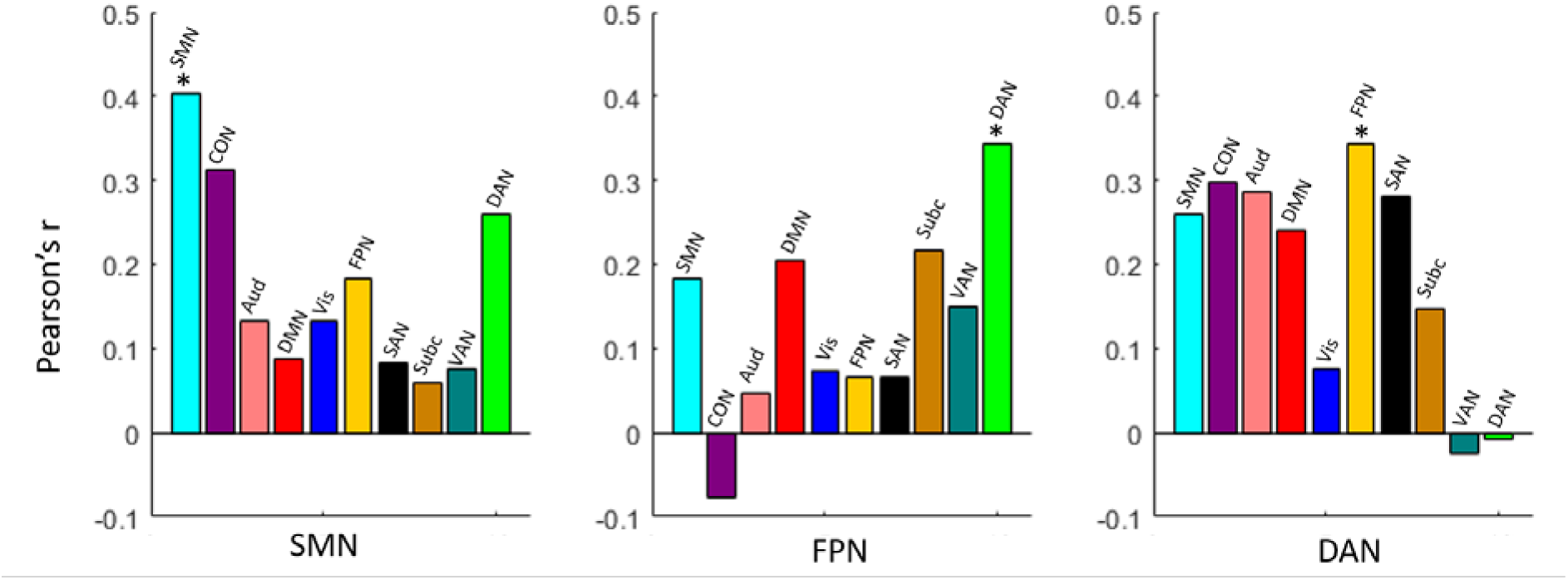
Multivariate pattern analysis results using within-network and between-network patterns. The x axis represents the three networks of interest (SMN, FPN, and DAN), and the y axis represents the correlation of observed and predicted EVET scores. Significant correlations of observed and predicted EVET scores were found using within-network connectivity patterns of the SMN and between-network connectivity patterns of FPN-DAN.

## 4. Discussion

The current results identified that functional connectivity patterns within the SMN and between FPN-DAN during rest predicted individual differences in multitasking ability using a virtual real-world task. Models based on these network patterns generalized to previously unseen individuals and could predict performance from resting-state connectivity alone. These results support previous research indicating that executive control regions in the frontal and parietal cortex underlie multitasking abilities (Al-Hashimi et al., 2015; Dux et al., 2009; Garner & Dux, 2015) as well as the role of the SMN in supporting fine-grained goal-directed action (Dixon et al., 2014; Dixon et al., 2017).

In recent years, connectome-based predictive modeling has been developed to predict cognitive measures such as sustained attention (Finn et al., 2015; Rosenberg et al., 2016; Shen et al., 2017). Although the functional organization of the brain is generally consistent across individuals, each person has their own unique pattern of functional connectivity, which can be used to predict individual differences in cognitive functioning. Furthermore, task performance can be predicted from functional connectivity patterns during resting-state (when participants are not engaged in any explicit task), which suggests that the neural architecture supporting cognitive ability is reflected in the brain’s intrinsic organization (Rosenberg et al., 2016). In influential models of large-scale functional networks that subserve goal-directed behavior, studies have identified networks consisting of the frontal and parietal cortices as a key component (Duncan, 2010, 2013; Fedorenko et al., 2013; Nelissen et al., 2013).

According to the framework of hierarchical organization of functional brain networks that underlie goal-directed action (Koechlin et al., 2003; Badre & D’Esposito., 2009), Dixon et al. (2014; 2017) suggested an anterior-to-posterior gradient consisting of the anterior FPN, posterior FPN, and SMN. The authors suggest that the anterior FPN is activated during the generation and monitoring of distal goals, whereas the posterior FPN is involved in guiding behavior toward proximal goals via executive control by encoding and maintaining task rules, and the SMN is involved in the execution of specific goal-directed actions. The DAN lies in between the FPN and SMN and is highly coupled and partially overlapping with the FPN during externally directed tasks (Dosenbach et al., 2008; Spreng et al., 2013; Yeo et al., 2011). The DAN is involved in top-down selection of behaviorally relevant stimuli (Corbetta & Shulman, 2002), and together with the FPN, is often commonly recruited during a variety of cognitively demanding tasks (Dosenbach et al., 2008; Fedorenko et al., 2013). This is in line with our finding that FPN-DAN connectivity is predictive of multitasking ability in individual participants. Once the FPN has established goals and task rules, the information needs to be translated into the execution of specific voluntary actions, which is established by the SMN (Bunge, 2004; Koechlin et al., 2003). The SMN is a set of highly interconnected somatosensory, primary motor, and premotor regions that interact to coordinate action. Within the SMN, the premotor and primary motor cortex operate in a hierarchy to translate visual and rule-based information into behaviorally appropriate motor responses (Bunge, 2004; Dixon et al., 2014; Rushworth et al., 2003). Here we show that within-network functional connectivity patterns in the SMN can predict how well an individual performs during naturalistic multitasking. Although we cannot test for hierarchical control in the current experiment, our findings support that the networks generally associated with various levels of cognitive control are related to individual differences in the ability to execute complex multi-component tasks.

As reviewed previously, multitasking is highly related to flexible cognitive control abilities. In particular, EVET taps into cognitive components including planning and execution, sustained attention, prospective memory, verbal and visuospatial working memory, task switching, processing speed, and cognitive bottlenecks, to provide a multifaceted measure of realistic everyday multitasking ability (Logie et al., 2011). We cannot separate multitasking from general fluid intelligence and cognitive control functions in this study as they highly overlap, but this would be an interesting questions for future studies, and may provide furture understanding of frontal lobe patients with intact intelligence but impaired multitasking skills (Shallice & Burgess, 1991; Burgess et al., 2000; Roca et al., 2011).

Previous studies have each identified parts of the FPN-DAN-SMN correlated with general cognitive or multitasking performance (e.g., Al-Hashimi et al., 2015; Cole et al., 2012; Erickson et al., 2007; Garner & Dux, 2015), yet all regions have been reported to be recruited more in multitask compared to single-task conditions during continuous multitasking (Al-Hashimi et al., 2015). Our experiment used a computer game to generate continuous engagement and a task with a set of goals, sub-tasks, and execution steps, a scenario that better reflects complex acts of real-word multitasking (Rothbart & Posner, 2015). Conventional multitasking paradigms used in the laboratory typically employ explicit instructions for participants to respond to each task separately even when they overlap in time (Al-Hashimi et al., 2015; Dux et al., 2009). However, real-world multitasking is more dynamic and allows more flexibility in scheduling parallel processes, which may differ from discrete task execution.

The use of a virtual task in this experiment may better emulate real-word multitasking and may explain why a large area of the prefrontal networks spanning the FPN-DAN-SMN may be related to individual differences in multitasking ability.

Connectivity of the frontal lobe and FPN have been implicated in cognitive control abilities in previous studies. However, our experiment did not show a significant relationship between global network connectivity and multitasking ability. A possibility that we did not observe significant relationships may be that we focused on global connectivity of the entire network, while regions within each network may have their unique characteristics. For example, Cole et al. (2012) found that global connectivity only in the LPFC, but not premotor cortex or medial posterior parietal cortex, showed correlation with fluid intelligence. When looking at global connectivity restricted to the LPFC, we find a significant relationship with multitasking performance. Other studies have found that although regions within the FPN is predominantly related to cognitive control and fluid intelligence, it is connectivity between distinct regions derived from multiple networks that together predict individual differences in these measures (Hearne et al., 2016; Song et al., 2008; Pamplona et al., 2015).

In univariate global connectivity analyses, a mean network summary measure is used to provide an average index of a network consisting of hundreds or thousands of connections to relate to individual differences (e.g., Cole et al., 2013; Cole et al., 2012; Rosenberg et al., 2016). Here in addition to the univariate global connectivity measure, we used a multivariate approach with SVR, thus taking all functional connections of a network into account. Such pattern information analysis can reveal effects lost in regional average analysis (Kriegeskorte & Bandettini, 2007; Mur et al., 2009). It is possible that using a virtual multitasking task assessed cognitive abilities that are not specific to one network, therefore diluting the effect after averaging. Yet, we were able to find strong effects with multivariate analysis, implying that the fine-grained connectivity patterns may be predictive of individual differences in multitasking ability, and can generalize to novel individuals (Dosenbach et al., 2010). Future studies with multivariate techniques in addition to conventional univariate analysis will likely shed light on how connectivity patterns are related to individual differences.

The current study used resting-state functional connectivity to predict multitasking performance outside the scanner. Recent studies have found that on top of the brain’s intrinsic connectivity, the functional network architectures can be reconfigured in response to an external task (Cole et al., 2014; Krienen et al., 2014). These changes are characterized by increases in functional connectivity strength as well as changes of selective patterns in line with increasing task demands (Cocchi et al., 2013; Hearne et al., 2017). Within the prefrontal cortex, the SMN transforms task rules and goal representations from the FPN into the execution of motor output (Koechlin et al, 2003; Dixon et al., 2017). Therefore, one might expect SMN-FPN or SMN-DAN connectivity during task to predict multitasking performance. Although we did not find this relationship with resting-state networks, it is possible that these relationships might emerge during task. While we cannot test this relationship in the current experiment, studying task-based connectivity with naturalistic stimuli would be an interesting avenue for future research.

In conclusion, our results show that resting-state functional connectivity patterns of networks involved in executive and sensorimotor control predict how well an individual performs in real-world multitasking. Previous studies suggest that these networks are organized to subserve goal-directed behavior at different processing levels. Individual differences in multitasking ability may result from how well these functional networks and cognitive components are orchestrated.

## Acknowledgements

We thank Yong-Quan Chen for help with data collection. We thank John Duncan for helpful comments and suggestions, and Yun-Hsuan Chang for help with data interpretation in the early stages of this study. We thank the Mind Research and Imaging Center (MRIC) at NCKU for consultation and instrument availability. This work was supported by the grants MOST 104-2420-H-006-004-MY2 and MOST 106-2420-H-006-005-MY2 from the Ministry of Science and Technology of Taiwan to SH. TW was supported by the Taiwan Cambridge Scholarship from the Cambridge Commonwealth, European & International Trust and the Percy Lander studentship from Downing College. The (Edinburgh Virtual Errands Task) EVET program was provided by Steven Trawley, Matthew Logie, and Robert Logie.

## References

Al-Hashimi, O., Zanto, T. P., & Gazzaley, A. (2015). Neural sources of performance decline during continuous multitasking. Cortex, 71, 49–57. doi: 10.1016/j.cortex.2015.06.001

Badre, D., & D’Esposito, M. (2009). Is the rostro-caudal axis of the frontal lobe hierarchical? Nature Reviews Neuroscience, 10, 659–669.

Bunge, S. A. (2004). How we use rules to select actions: a review of evidence from cognitive neuroscience. Cogn Affect Behav Neurosci, 4(4), 564–579.

Burgess, P. W. (2000). Real-world multitasking from a cognitive neuroscience perspective. Control of cognitive processes: Attention and performance XVIII, 465–472.

Cattell, R. B. (1967). The theory of fluid and crystallized general intelligence checked at the 5- 6 year-old level. Br J Educ Psychol, 37(2), 209–224.

Chen, T., Cai, W., Ryali, S., Supekar, K., & Menon, V. (2016). Distinct Global Brain Dynamics and Spatiotemporal Organization of the Salience Network. PLoS Biol, 14(6), e1002469. doi: 10.1371/journal.pbio.1002469

Cocchi, L., Halford, G. S., Zalesky, A., Harding, I. H., Ramm, B. J., Cutmore, T., … & Mattingley, J. B. (2013). Complexity in relational processing predicts changes in functional brain network dynamics. Cerebral Cortex, 24(9), 2283–2296.

Cole, M. W., Bassett, D. S., Power, J. D., Braver, T. S., & Petersen, S. E. (2014). Intrinsic and task-evoked network architectures of the human brain. Neuron, 83(1), 238–251.

Cole, M. W., Ito, T., & Braver, T. S. (2015). Lateral prefrontal cortex contributes to fluid intelligence through multinetwork connectivity. Brain connectivity, 5(8), 497–504.

Cole, M. W., Pathak, S., & Schneider, W. (2010). Identifying the brain’s most globally connected regions. Neuroimage, 49(4), 3132–3148.

Cole, M. W., Anticevic, A., Repovs, G., & Barch, D. (2011). Variable global dysconnectivity and individual differences in schizophrenia. Biol Psychiatry, 70(1), 43–50. doi: 10.1016/j.biopsych.2011.02.010

Cole, M. W., Reynolds, J. R., Power, J. D., Repovs, G., Anticevic, A., & Braver, T. S. (2013). Multi-task connectivity reveals flexible hubs for adaptive task control. Nature Neuroscience, 16(9), 1348–1355. doi: 10.1038/nn.3470

Cole, M. W., & Schneider, W. (2007). The cognitive control network: Integrated cortical regions with dissociable functions. Neuroimage, 37(1), 343–360. doi: 10.1016/j.neuroimage.2007.03.071

Cole, M. W., Yarkoni, T., Repovs, G., Anticevic, A., & Braver, T. S. (2012). Global connectivity of prefrontal cortex predicts cognitive control and intelligence. J Neurosci, 32(26), 8988–8999. doi: 10.1523/JNEUROSCI.0536-12.2012

Corbetta, M., & Shulman, G. L. (2002). Control of goal-directed and stimulus-driven attention in the brain. Nat Rev Neurosci, 3(3), 201–215. doi: 10.1038/nrn755

Dixon, M. L., Fox, K. C., & Christoff, K. (2014). Evidence for rostro-caudal functional organization in multiple brain areas related to goal-directed behavior. Brain Res, 1572, 26–39. doi: 10.1016/j.brainres.2014.05.012

Dixon, M. L., Girn, M., & Christoff, K. (2017). Hierarchical organization of frontoparietal control networks underlying goal-directed behavior. In The Prefrontal Cortex as an Executive, Emotional, and Social Brain (pp. 133–148). Springer Japan.

Dosenbach, N. U., Fair, D. A., Cohen, A. L., Schlaggar, B. L., & Petersen, S. E. (2008). A dual-networks architecture of top-down control. Trends Cogn Sci, 12(3), 99–105. doi: 10.1016/j.tics.2008.01.001

Dosenbach, N. U., Fair, D. A., Miezin, F. M., Cohen, A. L., Wenger, K. K., Dosenbach, R. A., … Petersen, S. E. (2007). Distinct brain networks for adaptive and stable task control in humans. Proc Natl Acad Sci U S A, 104(26), 11073–11078. doi: 10.1073/pnas.0704320104

Dosenbach, N. U., Nardos, B., Cohen, A. L., Fair, D. A., Power, J. D., Church, J. A., … Schlaggar, B. L. (2010). Prediction of individual brain maturity using fMRI. Science, 329(5997), 1358–1361. doi: 10.1126/science.1194144

Duncan, J.(2010). The multiple-demand (MD) system of the primate brain: mental programs for intelligent behaviour. Trends Cogn Sci, 14(4), 172–179. doi:10.1016/j.tics.2010.01.004

Duncan, J. (2013). The structure of cognition: attentional episodes in mind and brain. Neuron, 80(1), 35–50. doi: 10.1016/j.neuron.2013.09.015

Duncan, J., & Owen, A. M. (2000). Common regions of the human frontal lobe recruited by diverse cognitive demands. Trends in Neurosciences, 23(10), 475–483.

Dux, P. E., Tombu, M. N., Harrison, S., Rogers, B. P., Tong, F., & Marois, R. (2009). Training improves multitasking performance by increasing the speed of information processing in human prefrontal cortex. Neuron, 63(1), 127–138. doi: 10.1016/j.neuron.2009.06.005

Erickson, K. I., Colcombe, S. J., Wadhwa, R., Bherer, L., Peterson, M. S., Scalf, P. E., … Kramer, A. F. (2007). Training-induced functional activation changes in dual-task processing: an FMRI study. Cereb Cortex, 17(1), 192–204. doi: 10.1093/cercor/bhj137

Finn, E. S., Shen, X., Scheinost, D., Rosenberg, M. D., Huang, J., Chun, M. M., … & Constable, R. T. (2015). Functional connectome fingerprinting: identifying individuals using patterns of brain connectivity. Nature neuroscience, 18(11), 1664–1671.

Fedorenko, E., Duncan, J., & Kanwisher, N. (2013). Broad domain generality in focal regions of frontal and parietal cortex. Proc Natl Acad Sci U S A, 110(41), 16616–16621. doi: 10.1073/pnas.1315235110

Garner, K. G., & Dux, P. E. (2015). Training conquers multitasking costs by dividing task representations in the frontoparietal-subcortical system. Proc Natl Acad Sci U S A, 112(46), 14372–14377. doi: 10.1073/pnas.1511423112

Hastie, T., Tibshirani, R., & Friedman, J. (2009). The elements of statistical learning. New York: Springer series in statistics.

Hearne, L. J., Cocchi, L., Zalesky, A., & Mattingley, J. B. (2017). Reconfiguration of Brain Network Architectures between Resting-State and Complexity-Dependent Cognitive Reasoning. Journal of Neuroscience, 37(35), 8399–8411.

Hearne, L. J., Mattingley, J. B., & Cocchi, L. (2016). Functional brain networks related to individual differences in human intelligence at rest. Scientific reports, 6, 32328.

Koechlin, E., Basso, G., Pietrini, P., Panzer, S., & Grafman, J. (1999). The role of the anterior prefrontal cortex in human cognition. Nature, 399(6732), 148–151. doi: 10.1038/20178

Koechlin, E., Ody, C., & Kouneiher, F. (2003). The architecture of cognitive control in the human prefrontal cortex. Science, 302(5648), 1181–1185. doi: 10.1126/science.1088545

Kriegeskorte, N., & Bandettini, P. (2007). Analyzing for information, not activation, to exploit high-resolution fMRI. Neuroimage, 38(4), 649–662. doi: 10.1016/j.neuroimage.2007.02.022

Krienen, F. M., Yeo, B. T., & Buckner, R. L. (2014). Reconfigurable task-dependent functional coupling modes cluster around a core functional architecture. Phil. Trans. R. Soc. B, 369(1653), 20130526.

Laird, A. R., Fox, P. M., Eickhoff, S. B., Turner, J. A., Ray, K. L., McKay, D. R., … Fox, P. T. (2011). Behavioral interpretations of intrinsic connectivity networks. J Cogn Neurosci, 23(12), 4022–4037. doi: 10.1162/jocn_a_00077

Logie, R., Law, A., Trawley, S., & Nissan, J. (2010). Multitasking, working memory and remembering intentions. Psychologica Belgica, 50(3–4).

Logie, R. H., Trawley, S., & Law, A. (2011). Multitasking: multiple, domain-specific cognitive functions in a virtual environment. Mem Cognit, 39(8), 1561–1574. doi: 10.3758/s13421-011-0120-1

Manly, T., Hawkins, K., Evans, J., Woldt, K., & Robertson, I. H. (2002). Rehabilitation of executive function: facilitation of effective goal management on complex tasks using periodic auditory alerts. Neuropsychologia, 40(3), 271–281.

Marti, S., King, J. R., & Dehaene, S. (2015). Time-Resolved Decoding of Two Processing Chains during Dual-Task Interference. Neuron, 88(6), 1297–1307. doi: 10.1016/j.neuron.2015.10.040

Miller, E. K., & Cohen, J. D. (2001). An integrative theory of prefrontal cortex function. Annu Rev Neurosci, 24, 167–202. doi: 10.1146/annurev.neuro.24.1.167

Mur, M., Bandettini, P. A., & Kriegeskorte, N. (2009). Revealing representational content with pattern-information fMRI--an introductory guide. Soc Cogn Affect Neurosci, 4(1), 101–109. doi: 10.1093/scan/nsn044

Nelissen, N., Stokes, M., Nobre, A. C., & Rushworth, M. F. (2013). Frontal and parietal cortical interactions with distributed visual representations during selective attention and action selection. J Neurosci, 33(42), 16443–16458. doi: 10.1523/JNEUROSCI.2625-13.2013

Norman, D. A., & Shallice, T. (1986). Attention to action. In Consciousness and self-regulation (pp. 1–18). Springer US.

Pamplona, G. S., Santos Neto, G. S., Rosset, S. R., Rogers, B. P., & Salmon, C. E. (2015). Analyzing the association between functional connectivity of the brain and intellectual performance. Frontiers in human neuroscience, 9, 61.

Power, J. D., Cohen, A. L., Nelson, S. M., Wig, G. S., Barnes, K. A., Church, J. A., … Petersen, S. E. (2011). Functional network organization of the human brain. Neuron, 72(4), 665–678. doi: 10.1016/j.neuron.2011.09.006

Raven, J. C., & John Hugh Court. (1998). Raven’s progressive matrices and vocabulary scales. Oxford, UK: Oxford Psychologists Press.

Roca, M., Parr, A., Thompson, R., Woolgar, A., Torralva, T., Antoun, N., … & Duncan, J. (2009). Executive function and fluid intelligence after frontal lobe lesions. Brain, 133(1), 234–247.

Roca, M., Torralva, T., Gleichgerrcht, E., Woolgar, A., Thompson, R., Duncan, J., & Manes, F. (2011). The role of Area 10 (BA10) in human multitasking and in social cognition: a lesion study. Neuropsychologia, 49(13), 3525–3531.

Roca, M., Manes, F., Chade, A., Gleichgerrcht, E., Gershanik, O., Arévalo, G. G., … & Duncan, J. (2012). The relationship between executive functions and fluid intelligence in Parkinson’s disease. Psychological medicine, 42(11), 2445–2452.

Rosenberg, M. D., Finn, E. S., Scheinost, D., Papademetris, X., Shen, X., Constable, R. T., & Chun, M. M. (2016). A neuromarker of sustained attention from whole-brain functional connectivity. Nature Neuroscience, 19(1), 165–171. doi: 10.1038/nn.4179

Rothbart, M. K., & Posner, M. I. (2015). The developing brain in a multitasking world. Dev Rev, 35, 42–63. doi: 10.1016/j.dr.2014.12.006

Rubinov, M., & Sporns, O. (2011). Weight-conserving characterization of complex functional brain networks. Neuroimage, 56(4), 2068–2079. doi: 10.1016/j.neuroimage.2011.03.069

Rushworth, M. F., Johansen-Berg, H., Gobel, S. M., & Devlin, J. T. (2003). The left parietal and premotor cortices: motor attention and selection. Neuroimage, 20 Suppl 1, S89–100.

Shallice, T., & Burgess, P. W. (1991). Deficits in strategy application following frontal lobe damage in man. Brain, 114 *(Pt 2)*, 727–741.

Shen, X., Finn, E. S., Scheinost, D., Rosenberg, M. D., Chun, M. M., Papademetris, X., & Constable, R. T. (2017). Using connectome-based predictive modeling to predict individual behavior from brain connectivity. Nat Protoc, 12(3), 506–518. doi: 10.1038/nprot.2016.178

Song, M., Zhou, Y., Li, J., Liu, Y., Tian, L., Yu, C., & Jiang, T. (2008). Brain spontaneous functional connectivity and intelligence. Neuroimage, 41(3), 1168–1176.

Spreng, R. N., Sepulcre, J., Turner, G. R., Stevens, W. D., & Schacter, D. L. (2013). Intrinsic architecture underlying the relations among the default, dorsal attention, and frontoparietal control networks of the human brain. J Cogn Neurosci, 25(1), 74–86. doi: 10.1162/jocn_a_00281

Trawley, S., Law, A.S., Logie, M.R. & Logie, R.H. (2013). Desktop virtual reality in psychological research: a case study using the Source 3D game engine. EVET Technical Report.

Trawley, S. L., Law, A. S., & Logie, R. H. (2011). Event-based prospective remembering in a virtual world. Q J Exp Psychol (Hove), 64(11), 2181–2193. doi: 10.1080/17470218.2011.584976

Tschentscher, N., Mitchell, D., & Duncan, J. (2017). Fluid Intelligence Predicts Novel Rule Implementation in a Distributed Frontoparietal Control Network. J Neurosci 37(18), 4841–4847. doi: 10.1523/JNEUROSCI.2478-16.2017

Vincent, J. L., Kahn, I., Snyder, A. Z., Raichle, M. E., & Buckner, R. L. (2008). Evidence for a frontoparietal control system revealed by intrinsic functional connectivity. Journal of Neurophysiology, 100(6), 3328–3342. doi: 10.1152/jn.90355.2008

Yeo, B. T., Krienen, F. M., Sepulcre, J., Sabuncu, M. R., Lashkari, D., Hollinshead, M., … Buckner, R. L. (2011). The organization of the human cerebral cortex estimated by intrinsic functional connectivity. Journal of Neurophysiology, 106(3), 1125–1165. doi: 10.1152/jn.00338.2011

